# Machine learning for cell classification and neighborhood analysis in glioma tissue

**DOI:** 10.1101/2021.02.26.433051

**Authors:** Leslie Solorzano, Lina Wik, Thomas Olsson Bontell, Yuyu Wang, Anna H. Klemm, Johan Öfverstedt, Asgeir S. Jakola, Arne Östman, Carolina Wählby

## Abstract

Multiplexed and spatially resolved single-cell analyses that intend to study tissue heterogeneity and cell organization invariably face as a first step the challenge of cell classification. Accuracy and reproducibility are important for the down-stream process of counting cells, quantifying cell-cell interactions, and extracting information on disease-specific localized cell niches. Novel staining techniques make it possible to visualize and quantify large numbers of cell-specific molecular markers in parallel. However, due to variations in sample handling and artefacts from staining and scanning, cells of the same type may present different marker profiles both within and across samples. We address multiplexed immunofluorescence data from tissue microarrays of low grade gliomas and present a methodology using two different machine learning architectures and features insensitive to illumination to perform cell classification. The fully automated cell classification provides a measure of confidence for the decision and requires a comparably small annotated dataset for training, which can be created using freely available tools. Using the proposed method, we reached an accuracy of 83.1% on cell classification without the need for standardization of samples. Using our confidence measure, cells with low-confidence classifications could be excluded, pushing the classification accuracy to 94.5%. Next, we used the cell classification results to search for cell niches with an unsupervised learning approach based on graph neural networks. We show that the approach can re-detect specialized tissue niches in previously published data, and that our proposed cell classification leads to niche definitions that may be relevant for sub-groups of glioma, if applied to larger datasets.

## 1 Introduction

Spatially resolved single cell analysis allows for cell characterization in tumor microenvironment (TME) studies, and can be used by itself or in combination with other methods to find subgroups of disease that are associated with distinct clinical outcomes [1]. From the numerous multiplex staining methodologies (nicely reviewed in [2]), we want to mention two fluorescence based platforms that have complementary characteristics: i) co-detection by indexing (CODEX) [3] and ii) multiplexed immunofluorescence (mIF). While technologies like CODEX allow the simultaneous study of up to fifty biomarkers, it is low in throughput as it requires repeated rounds of staining and de-staining. In contrast, while mIF allows for the simultaneous staining of only up to eight markers, its high throughput nature allows to scan and analyze more samples. Previously, mIF has been used to quantify marker expression [4, 5, 6, 7, 8], and in conjunction with other techniques like flow cytometry, Hematoxylin and Eosin (H&E) staining, Immunohistochemistry (IHC) and single-cell RNAseq [9, 7, 10]. In [11] mIF stained cell populations in the colorectal cancer tumour microenvironment are identified using QuPath[12]. In [13] spectral unmixing of mIF stained cells is done using the InForm software and then classified based on manual thresholding and Random Forests (RF) using the HALO Highplex software. All these approaches include a manual adjustment of thresholds or normalization on a per-image basis, risking the introduction of bias. In [14] a more advanced pipeline is applied, including cell classification using a one-vs-all multi-class binary classification with support vector machines using InForm and Matlab.

To classify cells and study cell niches in tumor tissue, commonly, cells are first identified by segmentation of cell nuclei, followed by dilation to approximate a cell boundary, as implemented in popular open source software such as CellProfiler [15] and QuPath [12]. Private software for image analysis in mIF includes Akoya’s InForm, HALO and Aperio [2]. Closed software has a disadvantage of limited possibilities for sharing and reproducing projects and results. On the other hand, QuPath is open, offers the possibility of using RF but does not yet allow deep learning classifiers natively. However it includes scripting capabilities which makes it possible to extract the outlines of segmented cells and work with feature extraction and cell classification using other tools.

Typically, the next step in cell classification is marker quantification. Per cell marker quantification composes a profile that can be used to assign a class to a cell from an expected list of classes associated to a known marker panel. Fluorescent stains can have some degree of spectral overlap and bleed-through and each sample may have intensity variability due to sample-, staining- and scanning artefacts such as uneven illumination. To approach these challenges, a typical workflow of cell classification using mIF involves manual/visual adjustment and/or thresholding of images or per-cell measurements to define which cells are positive and negative for each specific marker. These adjustments are often made for each image and stain separately, making it a tedious process that is difficult to reproduce and may introduce bias. Stain normalization in histology is an intensely studied area [16], but few solutions work across large and multiplexed datasets where samples come from multiple sources, and the task becomes more challenging as more stains are added.

We present a methodology for fully automated cell classification that is able to provide a confidence measure, and compare two very popular and established methods in Machine Learning: i) an ensemble of fully connected neural networks (FCNNs) [17] and ii) Extreme Gradient Boosting (XGBoost) [18]. FCNNs create a weighted linear combination of the features which is able to take into account the variation within a class. Each of the neurons in each layer is connected to the adjacent layers and in this way all the knowledge is propagated. XGBoost is an advanced version of RF, an ensemble of decision trees, which, at each round of training, learns to focus on the hardest points to classify. Both methods use gradient information to arrive to a solution. But they are also very different, create different solution landscapes and have very different parameters and behaviors. We additionally explore using ensembles of classifiers, and propose a set of new marker intensity features that have less variation under uneven illumination, improving classification accuracy.

Following cell classification we proceeded to use the results for spatial analysis. Several methods and tools for exploration of spatial distribution of cells have been presented before [19, 20]. These tools are however focused on evaluating interactions among cell pairs, and not on niche discovery. With a small number of cell types, they could function also for niche discovery, but as cell number increases, pairwise comparisons will be limited. Instead, we approach cell niche characterization with an unsupervised graph neural network approach, spage2vec [21], originally developed for characterization of spatially resolved gene expression. We present a brief validation of its use on previously published spatially resolved single cell data obtained by CODEX [3] where we can compare to other niche detection approaches. Finally, we apply spage2vec to our TMA cores creating maps of local tissue niches with similar cellular neighborhoods.

## 2 Methods

### 2.1 Tissue collection and multiplexed immunofluorescence

In this study, we focus on primary brain tumors of the mutIDH1+ tumor subgroup. Primary brain tumors have, in general, been shown to be composed of variable tumor microenvironments, with demonstrated intra-case variability regarding both abundance and spatial organization of different cell types [22, 23]. In this setting the mutIDH1+ tumor sub-group presents special opportunities for cell profiling since the expression of the mutIDH protein in malignant cells allows distinction between malignant cells and astrocytes, which is difficult in many other primary brain tumors.

Lower grade gliomas were verified histopathologically, operated and embedded in paraffin between 2007 and 2016 at the Department of Neurosurgery, Sahlgrenska University Hospital, Gothenburg, Sweden. All procedures involving human specimens in this study were approved by the regional ethical committee in Gothenburg (Ep Dnr: 1067-16), and informed consent was given by the patients. Diagnoses were retrospectively confirmed and corrected by molecular classification. Tissue microarrays (TMAs) of tumor cores with a diameter of 1.2 mm were constructed at the Human Protein Atlas, Department of Immunology, Genetics and Pathology, Uppsala University, Sweden. The full cohort consists of two cores for each of 197 patients (394 cores in total) spread across four TMAs (four separate glass slides). Three of the TMAs are completely filled and contain 120 cores each while the last TMA contains 34 cores. Each TMA also contains two control cores from each of white and grey matter.

TMAs underwent octaplex immunofluorescence staining using an 7-plex Opal kit and additional Opal 480 and Opal 780 reagent kits (Akoya Biosciences, Marlborough, US) according to the manufacturer’s instructions. Briefly, Heat Induced Epitope Retrieval (HIER) was performed at pH6 for 5 minutes in a pressure cooker initially at 110 degrees, and at pH6 and 95 degrees in subsequent staining cycles. Akoya’s ready to use blocking buffer and secondary anti-mouse and –rabbit antibodies were incubated for 10 minutes. For Iba1, a separate biotinylated secondary antibody was used together with Vectastain Elite ABC kit (Vector Laboratories). Slides were washed three times in TBST between each step, and all Opal staining was developed for 10 minutes. Biomarkers selected for the study were Iba1, Ki-67, TMEM119, Neuro-Chrom a pan neuronal antibody coctail against NeuN, betaIII-tubulin, NF-H, MAP2 (a pan neuronal antibody cocktail against NeuN, beta III-tubulin, and NF-H, from now on referred to as NeuroC), MBP, mutIDH1, CD34, GFAP. Nuclei were counterstained with spectral DAPI provided with the kit for a 5-minute incubation at room temperature. TMAs were coverslipped using Prolong Gold antifade mountant (Thermo Fisher Scientific). For information on antibodies and specific staining conditions, see table S1 in supplementary material. Images were obtained after scanning TMAs through Vectra Polaris (Akoya Biosciences) at 20X magnification with optimized exposure time lengths, mapping each core with Phenochart image analysis software (Akoya Biosciences) and performing spectral unmixing using an in-house developed spectral library. Autofluorescence was subtracted using an image of unstained tissue exposed to corresponding HIER cycles as stained slides. Prior to cell classification, core edges as well as tissue artifacts with poor marker quality were masked away.

### 2.2 Ground truth for model training and evaluation

To create ground truth for model training and method evaluation we selected 21 random cores from the cohort of TMAs described above. The 21 cores were all from different cases. A higher number of training and evaluation samples would have required more manual annotations. Manual annotations are tedious, especially if multiple experts are involved. We concluded that 21 was a good balance between time investment and amount of data available for training and evaluation. Then, based on the mean marker intensity in each cell, extracted using QuPath as described above and shown in Figure S2, one person (Experts 1) manually selected a single threshold per marker and per image to indicate which cells in the image express the marker.

Cell type identification is not straight forward, and often relies on more than one biomarker per cell type. Additionally, several cell types may express overlapping markers. Therefore, we set up a classification scheme using a lookup table in order to systematically classify each cell based on the combined positivity/negativity for the markers in our biomarker panel. Rules for classification were based on existing knowledge about marker expression. Markers for astrocytes (GFAP), neurons (NeuN, betaIII-tubulin, NF-H, MAP2), myeloid derived macrophages and microglia (Iba1), endothelium (CD34), IDH1 mutated gliomas (IDH1 R132H) and proliferation (Ki-67) are well established. TMEM119 was recently shown to be specifically expressed in brain resident microglia and was used to distinguish between these cells and myeloid derived macrophages [24]. Due to the inability of the MBP marker to effectively stain oligodendrocyte soma, this marker turned out unsuitable for cell classification and was omitted from the analysis. It is also important to note that we included Ki-67 as a cell marker in our automated classification scheme although it is not cell-type specific, but rather a marker of cell proliferation. In the niche discovery explored here, this has little influence, but Ki-67 should be excluded from cell classification if proliferation is studied independently of cell type. A lookup table relating markers to cell type is shown in Figure S3.

We then proceeded to use the lookup table to assign a definitive training label based on the combination of markers in each cell. As a result, each cell received a single class label corresponding to one of six cell classes expected to be in the tissue: Astrocyte, Glioma, Neuron, Microglia, Macrophage, Endothelial. A small group of cells had combinations of markers that could not be adequately mapped to an expected cell type, and another sub group did not surpass any threshold resulting in no class. These two subgroups were deemed as “ambiguous” and were not used for training or evaluating the final automated classification results.

The subset of 21 cores labeled by expert 1 was further divided into two subsets, 11 cores for training and parameter optimization, and 10 cores for independent testing, taking care that the proportions of each cell class was as similar as possible in the two subsets. Due to class imbalance (as discussed later) we created subsets avoidng having zero cells representing a class in either testing or training. We made sure that cores were completely separate, none of the annotated cells used for training appears in the test set. Manual/visual determination of thresholds is challenging and highly subjective. We therefore let a different person (later referred to as Expert 2) independently determine thresholds for the 10 cores used for method evaluation. In total, Expert 1 classified 54953 cells from 21 cores, while leaving 8280 (15%) as ambiguous, out of which 1716 did not cross a threshold leaving them without a class, while the rest (6465 cells) had an unexpected combination of labels that did not match the predefined cell types. The large number of unexpected cells are likely mainly because of unspecificity of markers and hard thresholds not being flexible enough for local variations in marker signal intensity. There could also be other cell types in the tissue that do not fall in to any of the predefined categories. All ambiguous cells (unexpected class and no class) were removed from training and evaluation as they have no ground-truth label. Expert 2 classified 22293 cells in 10 cores, leaving 7992 (35%) as ambiguous.

### 2.3 Comparison with cell classification by average marker thresholds

As benchmarking for automating cell classification, we averaged all the manual thresholds for each marker across cores. We then applied these fixed thresholds to obtain a simple automated classification of the cells in the test set. This experiment functioned as a benchmark for comparison to the more advanced automated classification approaches described below.

### 2.4 Cell segmentation and feature extraction

Using QuPath and the DAPI channel as input, we segmented the cell nuclei in each core. Then we used dilation to define an area around each nucleus representing a region for quantifying cytoplasmic markers. It should be noted that, although we here on refer to this as a cell segmentation, this is in fact not a proper cell segmentation, which would be far more complicated due to the high variability of cell morphology in the tissue. All segmentation parameters were kept fixed for the full experiment as well as for creating ground truth.

After performing cell segmentation we quantified basic statistics of intensities (mean, min, max, and standard deviation) using QuPath. These features may however be influenced by errors in cell segmentation, e.g., small parts of neighboring cells reaching inside the very approximate cell outline may heavily influence a measure of mean intensity. They are also dependent on core-to-core variations in intensity.

We therefore exported the cell outlines and created a separate pipeline for further feature extraction and processing using Python Pandas library for data treatment and Scikit-Image [25] for image analysis. Using this pipeline we first measured the 90th percentile of intensity per cell and marker. Our reason for choosing the 90th percentile was to avoid noise from markers in neighboring cells. The 90th percentile also includes more information than the median (50th percentile) which would be close to zero when the stain is present only in the outer rim of the object, and not in the nucleus. With the 90th percentile we include more of the relevant information provided by the markers in the cell.

Next, we wanted to define a set of cell features with reduced core-to-core variation. While the intensities per marker and core may vary, the difference between markers remains similar from core to core. Hypothesizing that per-marker differences provide a valuable feature to circumvent core normalization we extracted the differences of 90th percentiles of intensity (here on referred to as d90s) for every pair of different markers. We reasoned that given that each class is determined by a combination of markers (as opposed to a single one), we provide the d90s to the models instead of them having to learn it. Leaving out DAPI and without repeating combinations of the remaining eight markers there are a total of 28 d90s. Concatenating these d90s to the regular intensity measurements: mean, minimum, maximum, standard deviation of the eight stains we obtain a feature vector per cell of length 68. We also experimented with other features from the intensities per cell such as the difference of median, mutual information and linear correlation (the former two being between pairs of markers inside the cell per pixel). These features were able to distinguish core similarities but they were not useful for a localized cell classification.

### 2.5 Machine learning models and cell classification confidence measures

#### 2.5.1 Data augmentation

Regardless of machine learning model, we augmented the training data to combat the imbalanced cell classes, with the glioma class containing about 100 times more cells than the other classes (Astrocyte: 1667, Glioma: 23168, Neuron: 695, Microglia: 230, Macrophage: 2498, Endothelial: 685). One could argue that a better balance may be obtained by including normal brain tissue samples. Such tissue is however not widely available, and typically has an architecture different from that of tumors. The augmentation procedure consisted of generating new “cell entries” by taking real cell feature values and adding Gaussian noise with zero mean and a standard deviation equal to each feature’s standard deviation. In this way every epoch of FCNN or XGBoost was trained with a new augmented training set which was created with a balanced number of cells per class (n=20000). Each model was given a set of cell labels and per-cell features for training. We compared two sets of features: i) the regular set of features per marker per cell (a total of 40 for ten channels (eight cell markers plus DAPI and autofluorescence) and four measurements: mean, min, max, and std dev) and ii) adding the 28 d90s, leading to a total of 68 features. The cell classes were Astrocyte, Glioma, Neuron, Microglia, Macrophage and Endothelial.

#### 2.5.2 Fully connected neural networks (FCNN)

We created a fully connected neural network (FCNN) architecture for cell classification consisting of an input layer with the size of the cell feature vector (size 68) and as output a feature vector of size six corresponding to the six different cell types. The model had three hidden fully connected layers in the middle, of sizes 100, 200 and 300 with rectified linear unit (ReLU) activations [26]. We used batch normalization [27] after the second and third hidden layer, before the activation functions, to reduce internal covariate shift and avoid convergence problems. To train the network, we used AdaBelief [28] as optimizer. This optimizer arrived to the solution twice as fast as a stochastic gradient descent optimizer [29] while providing similar classification accuracy. We trained each FCNN for 15 epochs.

#### 2.5.3 FCNN ensemble and confidence measure

Ensembles of machine learning models are well known to improve accuracy and robustness [30, 31]. To answer the question of whether an ensemble is needed and its advantages over a single model with the lowest (best) training loss, we ran a bootstrapping experiment and trained 100 unique models. Each new model was trained with a new seed, and each epoch was trained with a different augmentation of the training set.

We further hypothesized that a surrogate measure of confidence for the assigned class could be approximated from the votes by the ensemble of models. For each cell, we selected the mode of the votes as the true class and the percentage of models that voted for that class as a measure of how confident the ensemble as a whole was of that particular prediction.

All networks were trained using PyTorch 1.7.1 [32].

#### 2.5.4 Extreme gradient boosting (XGBoost)

For XGBoost [18] we used most of the default parameters, of note, seed per iteration True, tree depth of 7, eta 0.1, regularization alpha 0, regularization lambda 1, learning rate 0.05, objective ‘multi:softprob’ and number of rounds 200. This resulted in an ensemble of 1200 trees (given by the product of the number of classes and the number of rounds) which are weak classifiers, each tree being more specialized on the difficult training examples than the previous trees.

In order to obtain a measure of confidence for the XGBoost method, analogously to the FCNNs, we require an ensemble of models. We observed that even though an XGBoost model consists of an ensemble of decision tree models, they are neither independent nor equally weighted and therefore their agreement is not suitable as a measure of confidence. Instead we trained 100 XGBoost models and once more used the mode of their votes as the assigned class and the percentage of votes for the assigned class as a measure of confidence.

All XGBoosts were trained using the XGBoost Python API implementation version 1.1.1.

#### 2.5.5 Model implementation

FCNNs and XGBoost are not limited to a single programming language or software; what we present is a methodology which can be implemented in any existing tool. We would also like to mention that QuPath has some alternatives for the initial labeling, instead of using several single measurement classifiers, RF can be used on cores individually. However, using RF inside the software does not allow the user to determine which features to use for final classification.

If customized code is used outside of any particular software then more freedom of exploration is gained at the cost of increased technical difficulty. For the creation of the FCNN ensemble and XGBoost we used the popular programming language Python and the Pytorch library for Deep Learning. To obtain the d90s features we also use Python’s specialized image processing library scikit-image and Numpy. The code is available at https://github.com/wahlby-lab/ML-celltype-neighbor. Please note that this code is not a generic pipeline, but rather an implementation as it applies to our data; to use it with new data it has to be adapted. Image data of the cores used in this manuscript will be shared upon reasonable requests.

### 2.6 Cell niche detection by spage2vec

Spage2vec [21] is a graph neural network-based approach that learns a manifold of local object neighborhoods. The method takes object coordinates and labels as input, meaning that we could directly use coordinates and cell classes as output from the above presented methods. The learning is unsupervised, meaning that no *a priori* information on expected neighborhoods was needed. To validate the approach for niche detection, we used the publicly available CODEX cell class data provided in [3]. This data comes from healthy and diseased mouse spleen, which is clearly structured into functional niches. The authors define niches (semi-manually) by clustering cells based on the cell types they have in their immediate neighborhoods, resulting in tissue compartments. We therefore calibrated spage2vec parameters on this dataset and compared the output from spage2vec to previously presented compartments.

Spage2vec allows for the use of several network layers of any chosen size. We used the same parameters for our data and for the previously published CODEX data: two layers of sizes 50 and 2. This means that the resulting output manifold is a set of points in 2D space. We used an attention aggregator which results in the placement of these 2D points in a circle. Each point is separated from the others based on its difference in neighborhood. Distances are measured in terms of angles and cosine distance. We set the minimum distance that connects any two cells to be the 90th percentile of all the distances between each nearest cell in the dataset. This resulted in neighborhoods ranging from 1 to 10 cells.

## 3 Results

### 3.1 Comparison of classification accuracy

To evaluate classification performance, we used the designated test subset of 10 cores manually annotated with labels by two expert annotators as described in the previous sections. We regard annotations by expert 1 as ground truth and compare accuracy, i.e., number of correctly labeled cells over the total number of labeled cells, to the fully automated classification results produced by the FCNN and XGBoost ensembles, as well classification results obtained by average thresholds. We also compare to classification results obtained by manual classification by a second person, referred to as expert 2, see Figure 1.

**Figure 1:**
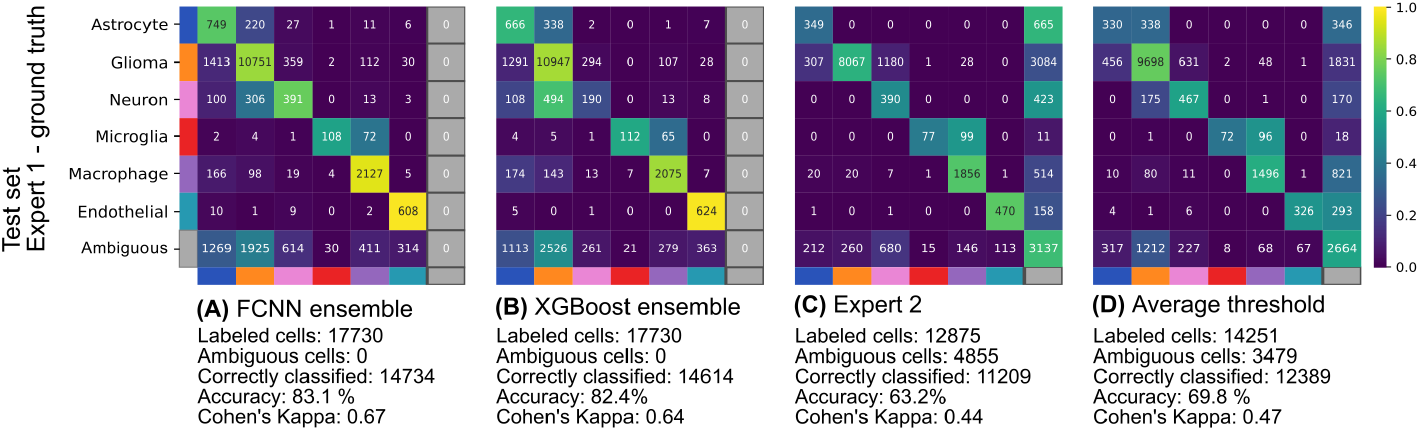
Confusion matrices comparing ground truth (as defined by expert 1) versus A) FCNN ensemble, B) XGBoost ensemble, C) human expert 2, and D) classification by average thresholds. The right-most column of each confusion matrix shows the number of cells not assigned to a class. The color coding in the confusion matrices reflect the proportion of the total number of cells in each class, including cells marked as ambiguous by expert 1. Accuracy measurements and Cohens Kappa are calculated only for cells in the test set that were assigned a class by expert 1.

When taking into account all the cells in the test set and using the full set of features including d90s, the FCNN ensemble reached 83.1% of accuracy while the XGBoost ensemble achieved 82.4% as compared to expert annotator 1. Classifying cells by applying averages of thresholds from the training set resulted in an accuracy of 69.8% on the test set (as compared to non-ambiguous cells in the ground truth by expert 1), and as many as 3479 out of the 17730 of the cells in the test set were left without an assigned class, as shown the confusion matrices in Figure 1. It can also be observed that the FCNN ensemble, as well as the XGBoost ensemble, assigns a class to every cell (leaving no ambiguous cells) while expert 2 as well as classification by averaged thresholds (1 C and D) result in many ambiguous cells, and thus reduced recall. The bright diagonals in both 1 A and B indicate that FCNNs and XGBoost ensembles have high accuracy and recall, but also that they have some difficulty separating glioma cells from neurons and microglia from macrophages.

### 3.2 Confidence thresholding increases precision at the cost of sensitivity

Using ensembles of classifiers allowed us to provide a measure of the confidence of the class voted by the ensemble. Note that all cells above the confidence threshold will be assigned a class, which may be a true (TP) or a false (FP) as compared to the class assigned by expert 1. Cells that fall below the confidence threshold are regarded as false negatives (FN). Figure 2A shows that applying an increasing confidence threshold increases the classification precision (TP/(TP+FP)), while at the same time decreasing sensitivity (TP/(TP+FN)). Thus, the confidence score can be used as a criterion to exclude ambiguous cells or to weight cells and their contributions to further downstream analysis of cell niches. In Figure 2A it can be observed that the FCNN ensemble achieved its highest precision value of 94.47% when the confidence threshold was set to the maximum (1.0) while the XGBoost ensemble reaches 88.3% meaning that it has less discriminative power, and regardless of the cost, no higher precision can be achieved.

**Figure 2:**
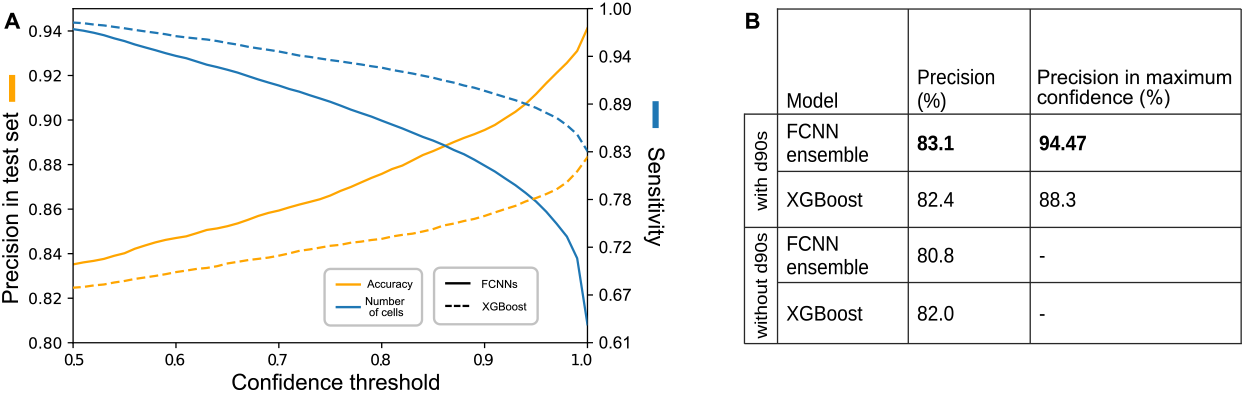
Confidence scores and effect of d90s features. A) Confidence in cell class, defined as the percentage of ensemble models that voted for the mode of the ensembles (x-axis), plotted against precision and sensitivity at varying cell class confidence thresholds. A high confidence thresholds leads to high precision (orange), at the cost of sensitivity (blue) for both models. B) Summary of precision achieved with and without d90s features and when using only maximum confidence.

### 3.3 Feature engineering

We studied the benefit of including the d90s features in relation to the performance of both machine learning models. We trained 100 components per model, one including the d90s and one that only included common marker intensity features. Each ensemble architecture used the same parameters for both sets of features. Both ensembles increased in precision when presented the d90s features. Without d90s, the FCNNs had a reduced precision of 80.75% and the XGBoost remained close at 82% (with only a drop of 0.4%). This showed that the d90s features helped the classification mostly for the FCNNs. Figure 2B summarizes the precision of each ensemble with and without d90s features.

### 3.4 Ensembles perform better than single models

FCNNs and XGBoost are very different architectures and even if they achieved similar performance they do so for different reasons. We explored how a single model arrives to a conclusion and what expected value of accuracy can be reached if we could train an infinite number of models.

Figure 3A shows the relationship between training loss and test set accuracy for each of 100 trained FCNNs. Using the annotated test set we could conclude that there was no correlation between training loss and test set accuracy, meaning that the FCNN with the lowest training loss can not be expected to perform well on the test set (correlation coefficient of −0.22). Additionally, the density of networks above the ensemble accuracy is very low, meaning that the likelihood of obtaining an FCNN that can outperform the ensemble is very low.

**Figure 3:**
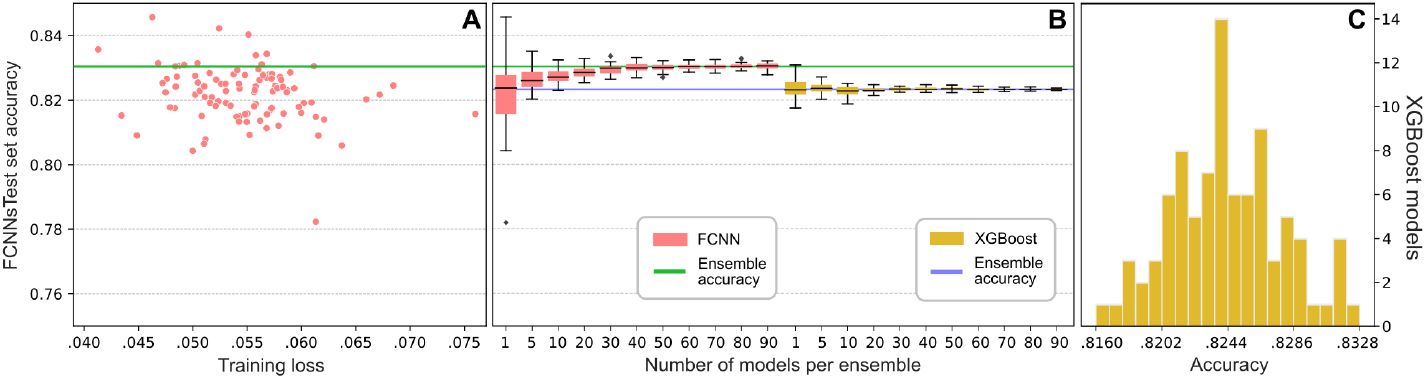
A) Scatterplot relating the training loss versus test accuracy of one hundred unique FCNNs trained to explore the solution landscape. B) Boostrap experiment to estimate the expected test accuracy using different amount of models in the ensembles. C) Histogram of accuracies amongst XGBoost models.

To estimate the effect of varying the number of models in the ensemble, we selected 30 random ensembles for each considered number of models (30 ensembles of sizes 1, 5, 10 etc) and quantified their accuracy, displaying the resulting accuracy distributions in Figure 3B. We observed that an ensemble of 30 FCNNs or more yields an expected value of 83.1% accuracy on the test set, and that increasing the number of networks further does not increase the accuracy by a large margin. For XGBoost the expected value does not change noticeably and remains around 82%. Based on training loss, XGBoost always achieved 0 loss so we cannot choose a model based on training loss, instead, we show in 3C a histogram of the number of models per accuracy. Low and high accuracies in the XGBoost range have lower probability of appearing so we can expect a total accuracy of 82% with little to no variability which means that almost all XGBoost models arrive to the same conclusions which is expected given the nature and limitations of the architecture.

### 3.5 Towards cell niche analysis using spage2vec

Spage2vec has previously only been used to find niches of gene expression, and to verify spage2vec as a method for detecting cell niches we performed an exploratory analysis and comparison with a previously study of tissue architecture from CODEX data of the mouse spleen [3]. Spage2vec offers several interesting features. It associates a value to a cell based on its neighborhood, meaning it characterizes each neighborhood, offering a measure of difference between neighborhoods. Additionally, since spage2vec in gene expression can be associated to spatial compartments and cell types, we wanted to observe if spage2vec applied to cell types within tissue would have a relation to spatial compartments, or niches. Supplementary Figure S4 shows the result of this analysis. The full dataset contains nine samples, all of which were used to train spage2vec. Figure S4 (A) shows the first 3 samples for which delineations of compartments were provided in the previous publication. (B) shows the cells color coded with spage2vec representations, correlating with compartments in (A). The semi-manually created maps in (A) are coarse and limited, while in (B) the representation is continuous and can be divided in several compartments and more fine grained information can be appreciated. (C) shows the amount of cells per spage2vec angle and peaks can be associated to regions with more homogenous cell neighborhoods. Note that colors that are present in (C) but not in (B) belong to the six samples not shown here, likely representing niches typical for diseased tissue. (D) shows a the spage2vec composition of each CODEX spatial compartment for further comparison. Although the size and type of niches found in in the CODEX dataset may not correspond to the size and type of niches one would expect in glioma tissue cores, it functioned as a proof-of-principle for general niche detection. The size of the resulting niches in the two datasets (despite using the same input parameters) also makes it clear that heterogeneity is higher in our TMA cores.

Having confirmed that spage2vec is applicable single cell data we proceeded to use the method and the output from our cell classification to find cell niches in our TMA cores. Resulting common and structured cell niches can be observed in Figure 4 (A), where similar colors correspond to similar niches. The angle histogram of the spage2vec values in Figure 4 (B) shows the frequency (x-axis) of different niches (y-axis), making it clear that some niches are more common than others (note log scale). Figure 4 (C) shows the cell-neighborhood composition of each niche, and the peaks in the spage2vec histogram coincide with peaks in the mean class representation in the niche histogram. A more detailed example of the cell composition of a given niche is shown in Figure S5. This indicates that spage2vec is able to describe a niche with a single value and is able to organize niches based on similarity, capturing distinct class compositions. High values in the mean cell composition indicate niche compositions with low standard deviation and therefore homogeneous niche cell composition, while low values indicate very heterogeneous niches.

**Figure 4:**
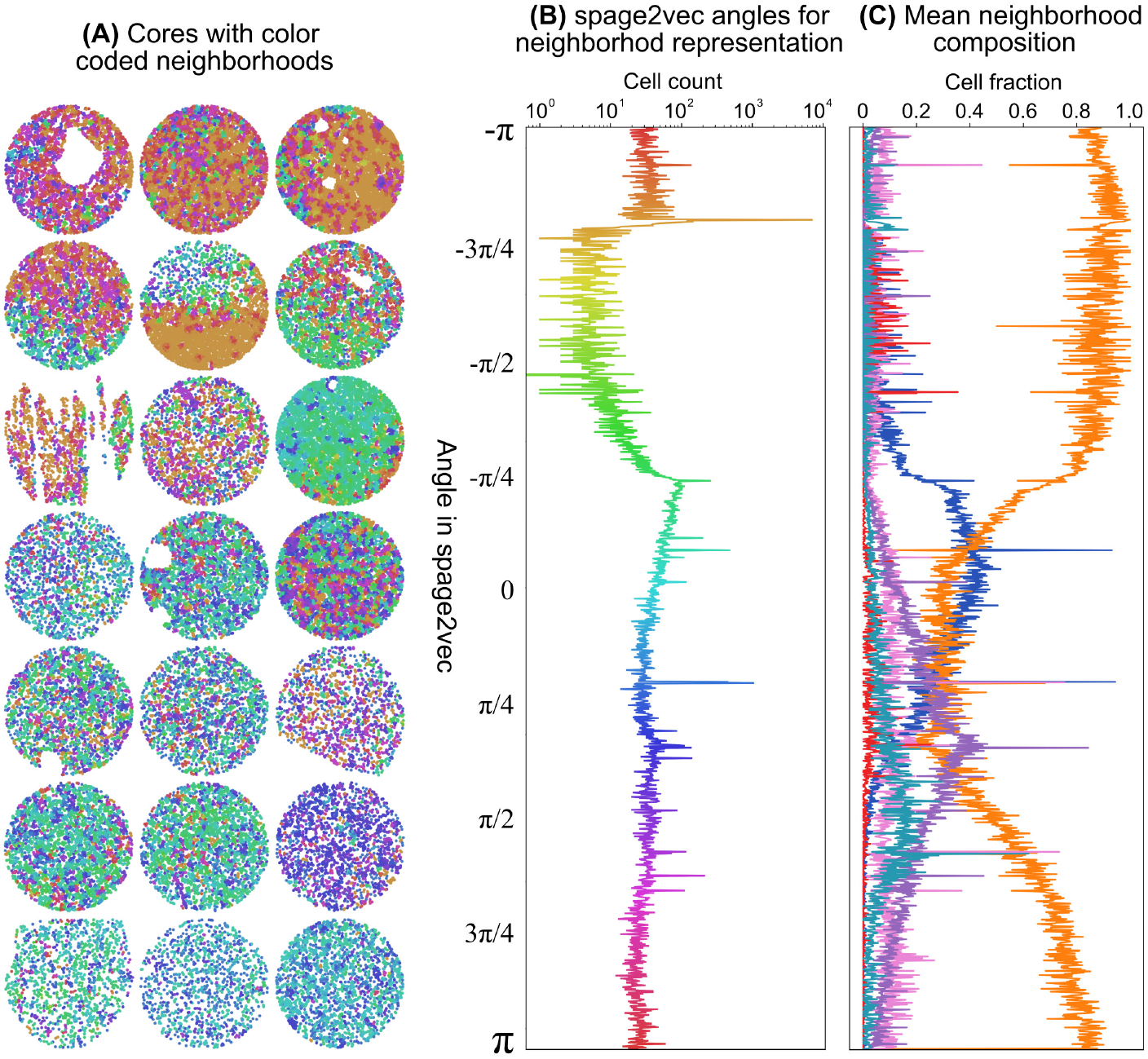
Cell niches defined by spage2vec on glioma TMA cores. A) Each cell is colored according to the niche it belongs to, with similar colors representing similar niches (represented by angle in B). The diameter of each core is approximately 1.2 mm. B) The cell count per niche (x-axis) is plotted against niche (y-axis) as a spage2vec angle histogram. With this representation of the spage2vec embedding, niches with similar cell composition are close in angle and color, while niches with very different composition are further apart, and peaks represent highly abundant niches. C) The cell composition per niche is presented as normalized mean niche composition in a bin centered in the same spage2vec angle. The colors are the same as in figures in the paper: Astrocyte (blue), Glioma (orange), Neuron (pink), Microglia (red), Macrophages (purple), and Endothelial (navygreen).

## 4 Conclusions and discussion

We have presented a methodology for cell classification using two popular machine learning architectures FCNN and XGBoost. Accurate cell classification is necessary for further analysis of tissue architecture and cell niches. We also presented the use of the spage2vec [21] method for the analysis of spatial cellular composition of tissue on the results from our classification.

We show that ensembles of classifiers arrive to similar conclusions as our two expert annotators which encourages us to continue using this method to classify larger cohorts of TMA cores and avoid manual thresholding. However, Figures 1, S6 and S7, show a confusion between the glioma class (orange) and neurons (pink). Confusion between microglia and macrophages is also noted. In order to study the possible cause we looked more carefully at some raw images. Figure S8 (A), and zoom-in (B), show a core where all outlined cells were labeled as glioma cells by expert 1. The cells with orange outlines were correctly classified as glioma cells by the FCNN ensemble, while those with orange outlines were misclassified as neurons. (C) show that the pink (incorrect) cells are distributed all through the core and their appearance does not seem to be due to imaging artefacts. Next, (D) and (E) show masked cells with a miniview of all independent markers, illustrating that it is very difficult to separate incorrectly classified glioma cells (D) from correctly classified glioma cells (E) by visual observation as all these cells are positive for both NeuroC (pink) and mutIDH1 (orange). Also quantitatively, differences are difficult to find; (F) compares average feature profiles for correctly and incorrectly classified glioma and neuron cells. Both glioma and neurons have a high value of the NeuroC marker, as highlighted in green, showing that the NeuroC marker is a source of confusion.

The same rationale goes for the profiles of macrophages and microglia, which are both IBA1 positive and their separation thus depends mostly on TMEM119 status, resulting in confusion between the classes as shown in figure S9. Note as well that these four cell types are also the cell types where concordance is lowest in the comparisons of the two expert annotators (Figure 1 C). The limitations of the automatic scoring are thus most likely due to the quality of the raw data, rather than poor performance of the FCNN ensemble. For continued studies, where the methods will be applied to address biomedical research questions, alternative markers should be considered.

High throughput image analysis of TMAs generally requires sample standardization or working on samples individually. The proposed methodology using ensembles is able to learn sample distributions, thus circumventing standardization difficulties. We presented the d90s features that capture information about relative cells marker profiles, leading to an increase in accuracy. From all XGBoost models, some of the d90s features are in the top 20 (out of 68) features in the measures of gain, coverage and weight (Figure S10), showing that they are indeed important features for decision making.

Furthermore, we showed that using an ensemble is better than using a single model and that the voting scheme can be converted to a measure of cell classification confidence. We also showed that within the highest confidence classifications there is indeed higher classification precision. This is especially useful if the set of markers does not cover all possible cell types that could be present in the tissue as confidence thresholding could potentially be used to flag cells that do not belong to the pre-assigned cell classes defined by the markers. If high precision is preferred at the cost of removing cells from analysis (i.e. lower sensitivity) then the confidence is a good parameter for thresholding. If a higher number of classified cells are preferred, the confidence threshold can be lowered and more cells are included. When the confidence is visualized as a colored map (supplementary FigureS11) we observe that the confidence is uniformly distributed indicating that our confidence measure is independent on whether a core belongs to training or testing sets and it does not follow any spurious pattern, increasing our trust on the map and the measure itself.

After comparing FCNN and XGBoost and exploring what both methods offer one could consider adapting strategies from both architectures. XGBoost gradually focuses on hard examples. A similar strategy could be used in FCNNs by implementing a weighting scheme where sample weights change as samples become easier or harder to classify. Also after feature exploration with XGBoost one could perhaps rethink the choice of features for the FCNN and possibly increase accuracy.

Finally we show that our cell classification allows us to perform further spatial analysis of the cell niches and allows us to create a grouping of TMA cores. Near future work includes using the proposed cell classification and spatial niche discovery to explore larger cohorts of TMA cores and relate them to clinical data.

## 5 Acknowledgements

Prof. Anja Smits, MD, PhD, Inst. of Neuroscience and Physiology, Dept. of Clinical Neuroscience, University of Gothenburg, Sahlgrenska Academy, Sweden, is acknowledged for facilitating the study and for contributing to creation of the tissue collection.

## 6 Funding

This project was financed by the European Research council via ERC (Ref 682810), the Swedish Foundation for Strategic Research (grant BD150008), the Swedish Research Council (Vetenskapsrådet) and the Swedish Cancer Society (Cancerfonden). Support was also provided by the SciLifeLab BioImage Informatics Facility.

## 7 Conflict of interest

The authors declare no conflict of interest

## 8 Supplementary

**Figure S1:**
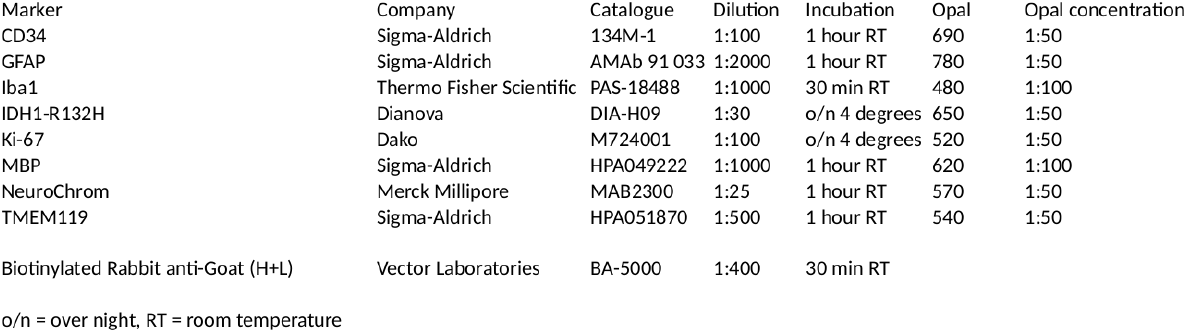
Table: Antibodies and specific staining conditions

**Figure S2:**
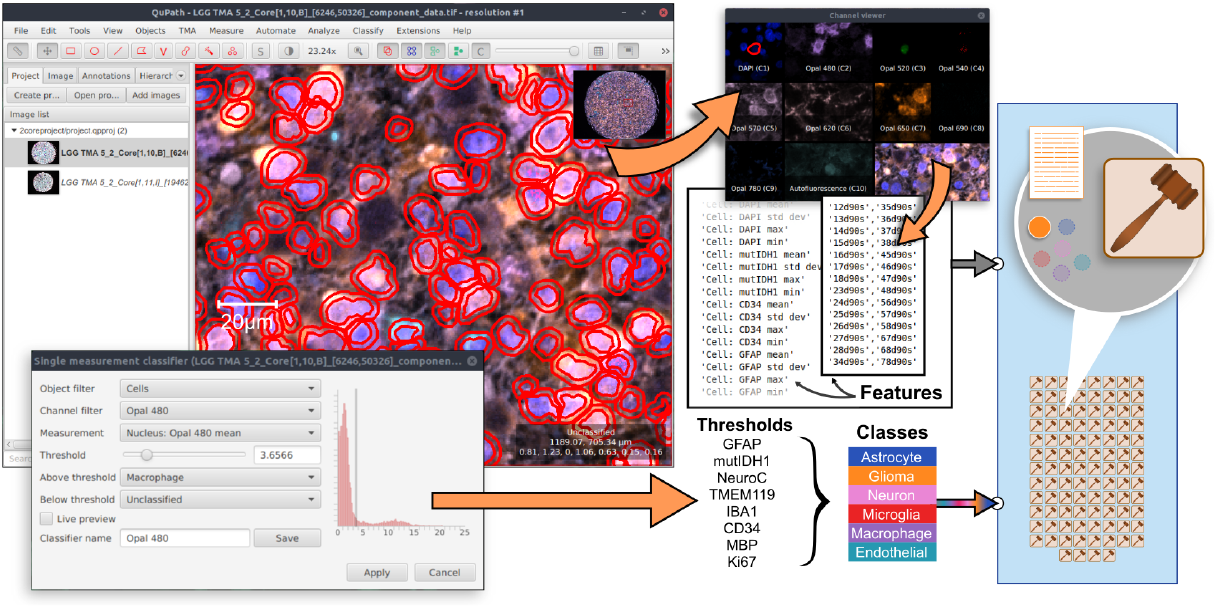
Creating training data: Example of the QuPath interface to perform cell segmentation of a single core and manually select thresholds per marker to generate cell classes. The cell segmentation masks are also exported to enable extraction of the d90s features outside of QuPath. These features are input to the FCNN and the XGBoost ensembles.

**Figure S3:**
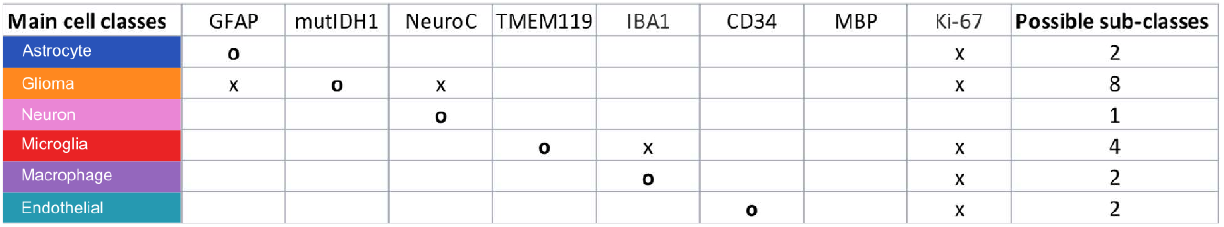
Marker to cell class lookup table, where “o” means main marker while “x” means possible additional marker. A cell’s marker intensity value has to cross the “o” marker threshold and may cross any “x” marker threshold to belong to the corresponding cell class. Cells whose marker intensities do not match the patterns in the table, or have values lower than all thresholds are considered “ambigous”.

**Figure S4:**
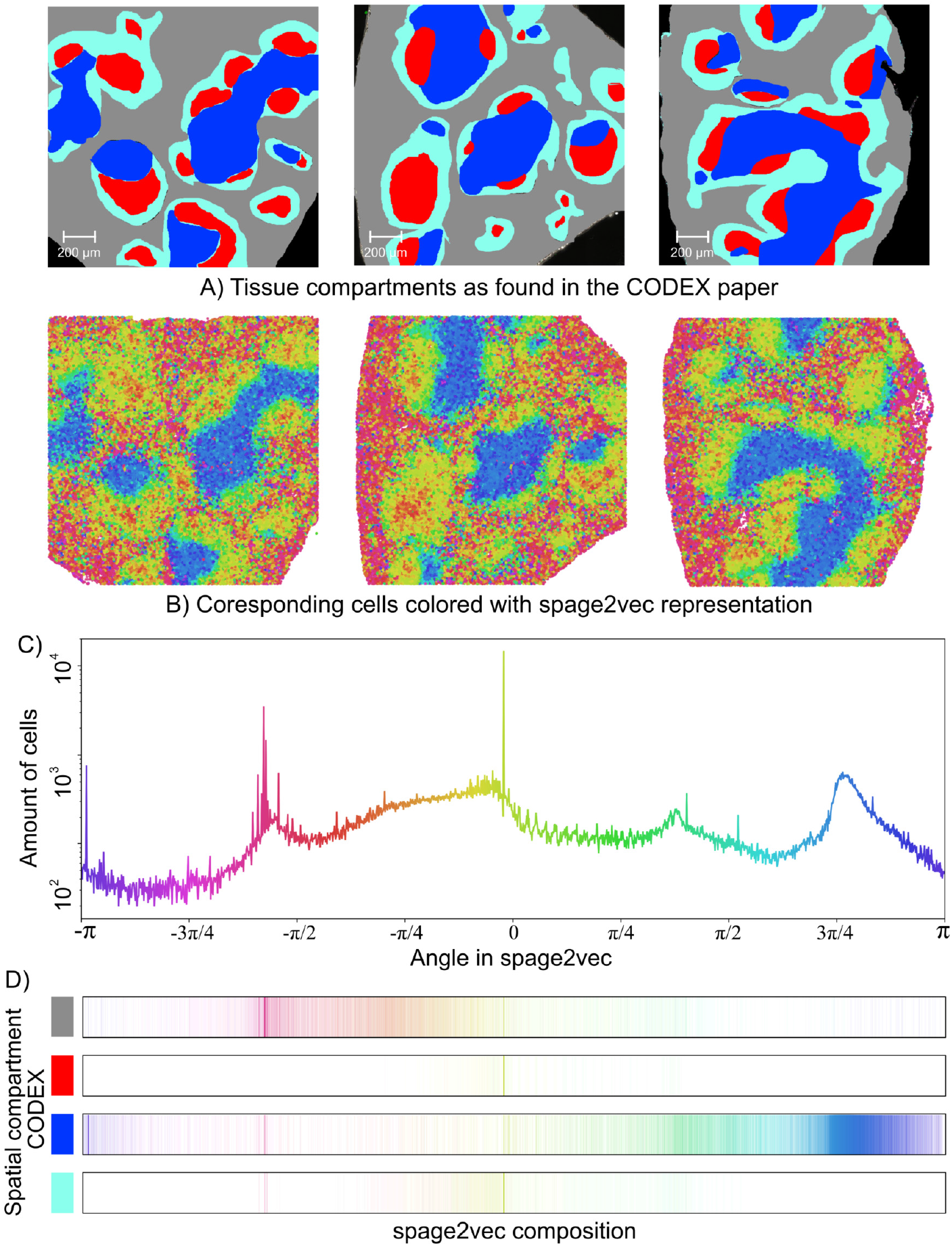
Niche detection by spage2vec applied to CODEX data from [3]. A) Semi-manually drawn spatial compartment maps of healthy mouse spleen provided in the previous publication. B) Cell niches as produced by applying spage2vec to all cells from a total of nine tissue samples presented in [3]. It can be observed that the neighborhood types are clustered in a similar way as in A. C) Shows the spage2vec histogram with the amount of cells per cell niche, represented by angle. D) shows the spage2vec niche composition for each of the of the compartments presented in (A). CODEX uses *a priori* knowledge to create the regions in a semi automatic way. However the maps are similar eventhough the goal of spage2vec is to characterize neighborhoods which do not necessarily have to match regions. This is a proof of concept to show that the neighborhoods description is not random. First row of tissue compartment images belongs to the authors of the CODEX publication in [3].

**Figure S5:**
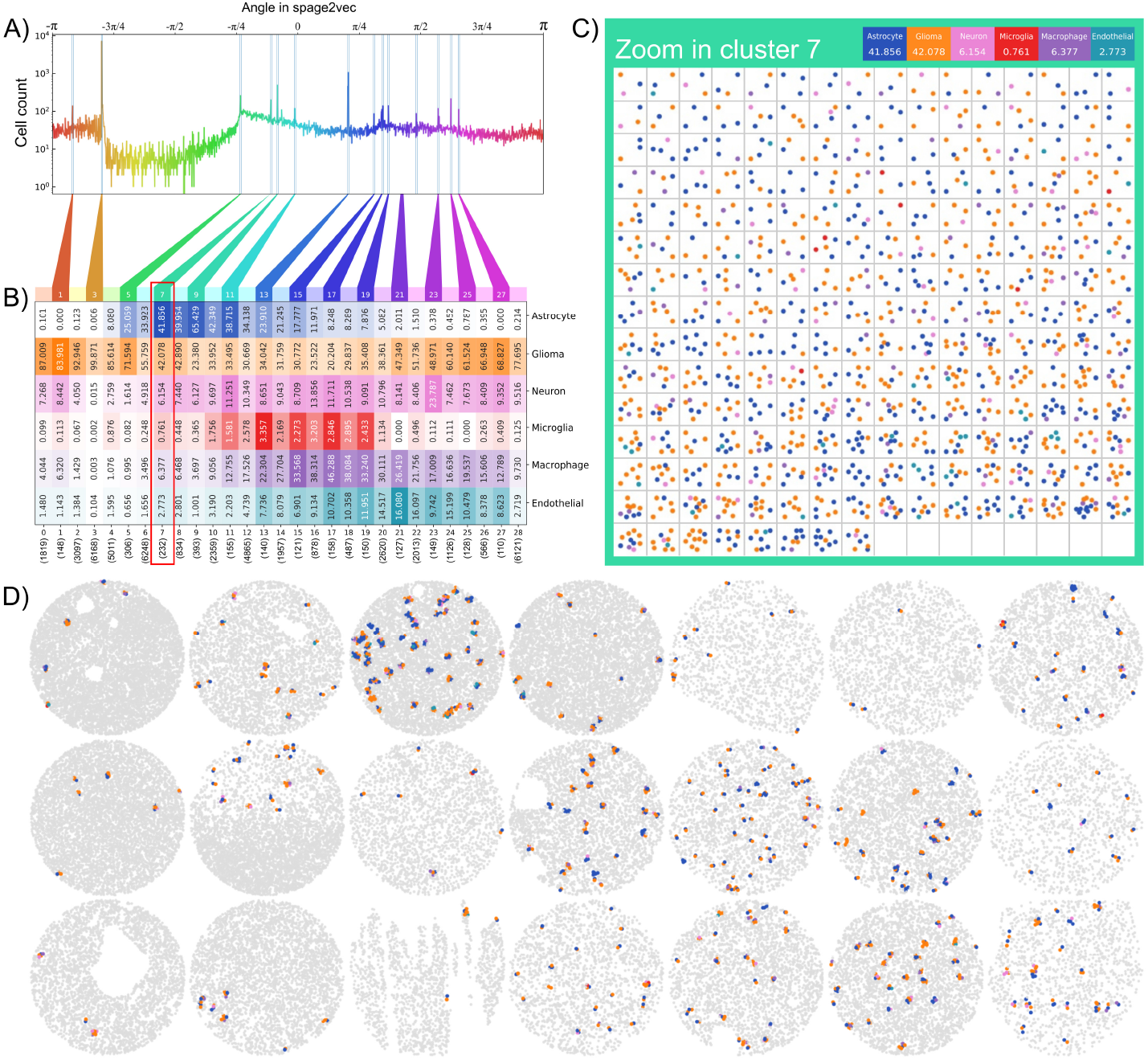
Spage2vec cluster zoom-in for glioma data. A) Histogram showing the amount of cells per angle in spage2vec. Peaks in the histogram are clusters (niches) that are common. B) Heatmap where he numbers in the columns mean the chance that any given neighborhood in this cluster contains cells of this class. The color strength reflects how many of the total available cells for this class in the dataset are present in this cluster. Although the numbers may correlate with the colors they mean different things. For example, there are very few endothelial cells in total (685 our of 54000) and most of them are in cluster 21. But if we look at each individual cluster, there is still a low probability (j16%) of finding an endothelial cell. C) Display of all the neighborhoods in cluster 7. These neighborhood type has 41% chance of having astrocytes, 42% chance of having glioma, 6% of having neurons, 6% of having macrophages, 2.7% of having endothelial and 0.7% of having microglia. Clusters are shown individually and in D) their spatial location in the cores. The diameter of each core is approximately 1.2 mm.

**Figure S6:**
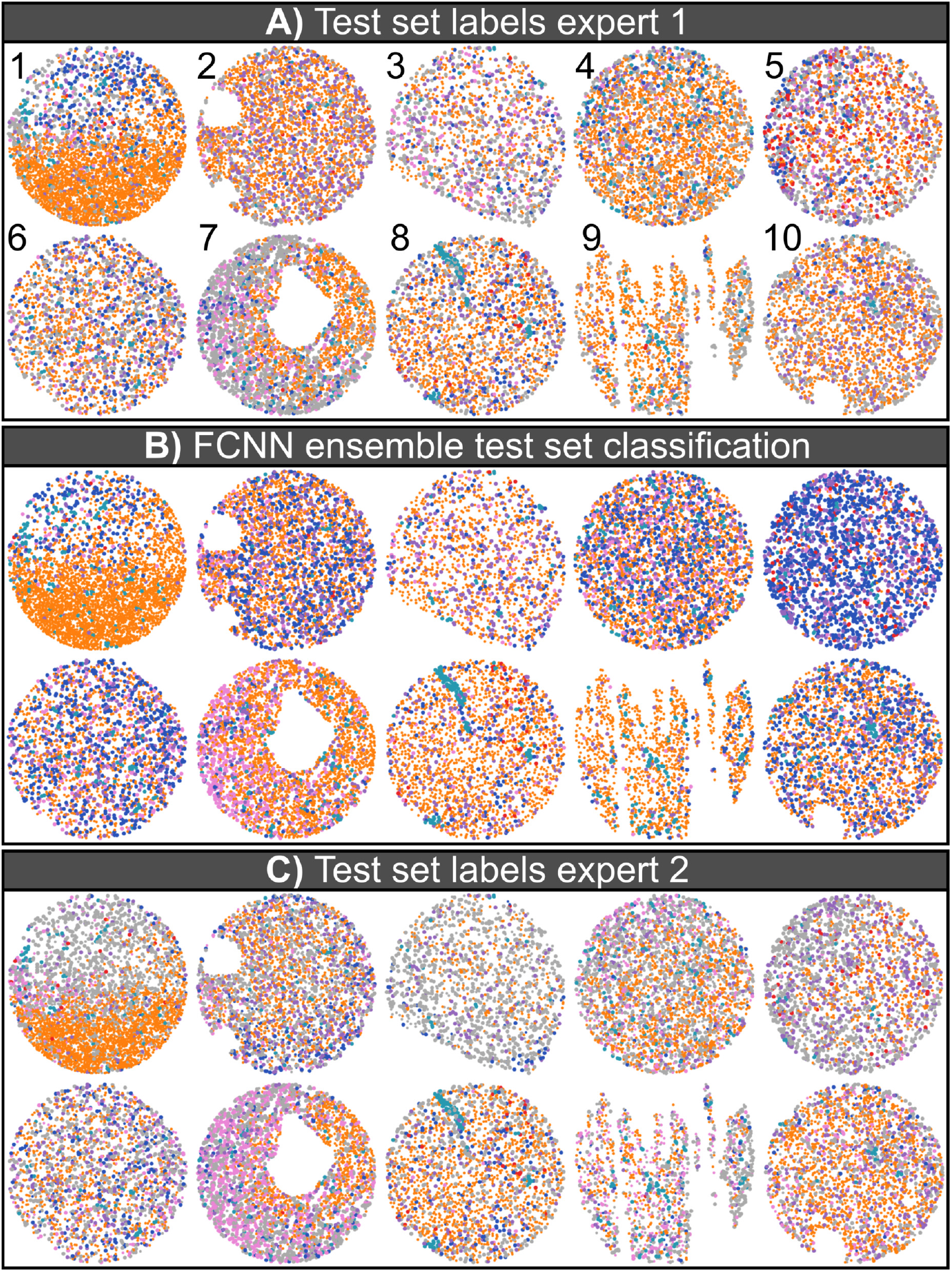
Cell classification results on the 10 independent test cores compared amongst experts and the FCNN ensemble. The figure serves to visualize agreements and differences amongst experts and the FNN ensemble. A) Cell classes from manual/visual annotation by expert 1. B) Cell classification results by the FCNN ensemble. C) Cell classes from manual/visual annotation by expert 2. Grey color means that the cell has no label due to ambiguity. More in depth comparison presented in the continuation of this figure in, figure S7. The diameter of each core is approximately 1.2 mm.

**Figure S7:**
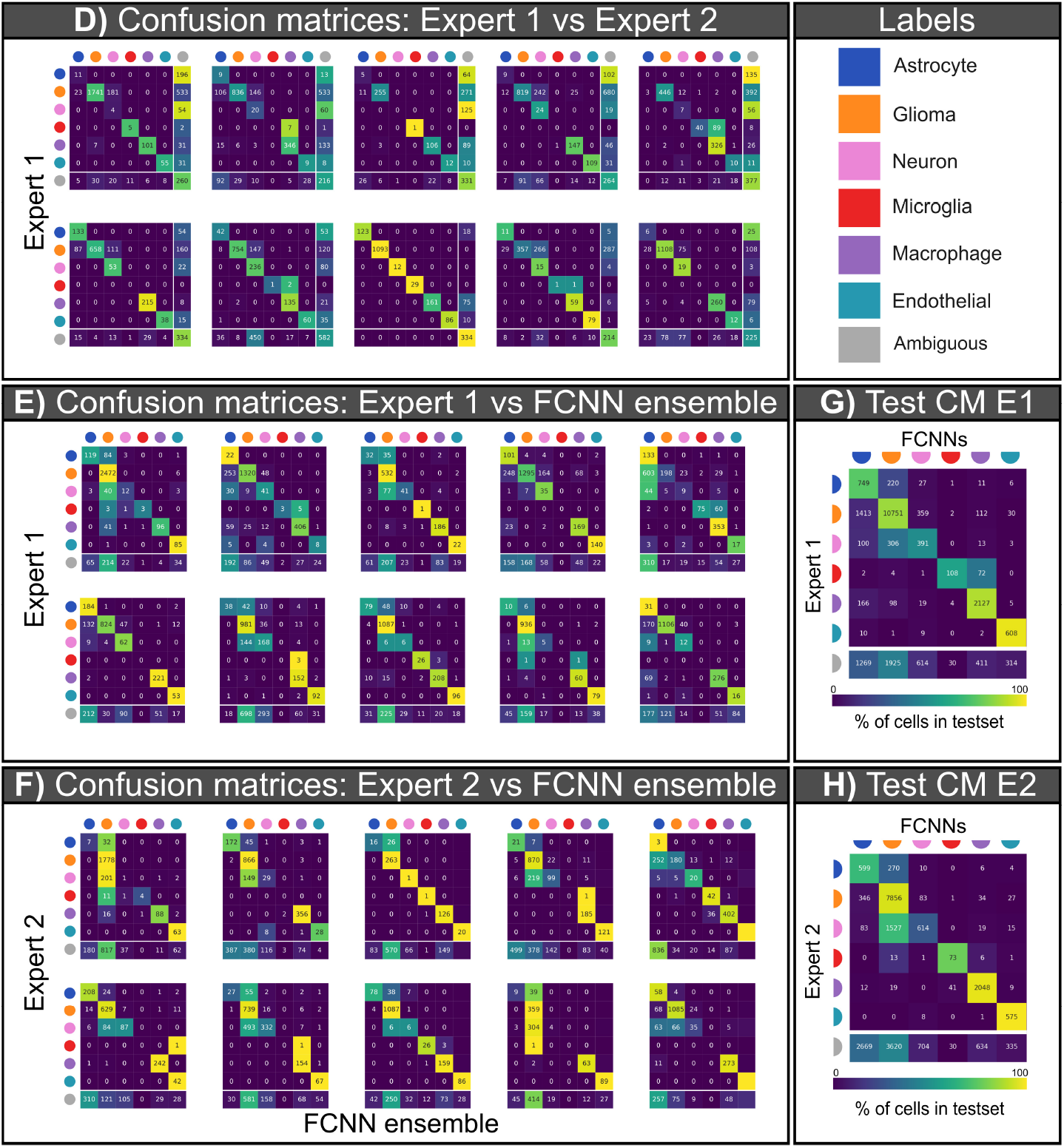
Confusion matrices showing agreements between experts and FNN ensemble corresponding to the cores in figure S6. D) Confusion matrices per core showing the agreement between expert 1 and expert 2. Notice the number of ambiguous cells provided in the bottom row in each confusion matrix, and how they reduce the number of agreed cells. E) Confusion matrices between expert 1 and the FCNN ensemble. F) Confusion matrices between expert 2 and the FCNN. G) Summary confusion matrix between expert 1 and the FCNN. H) Summary confusion matrix between expert 2 and the FCNN. D, E and F correspond to A, B and C in figure S6

**Figure S8:**
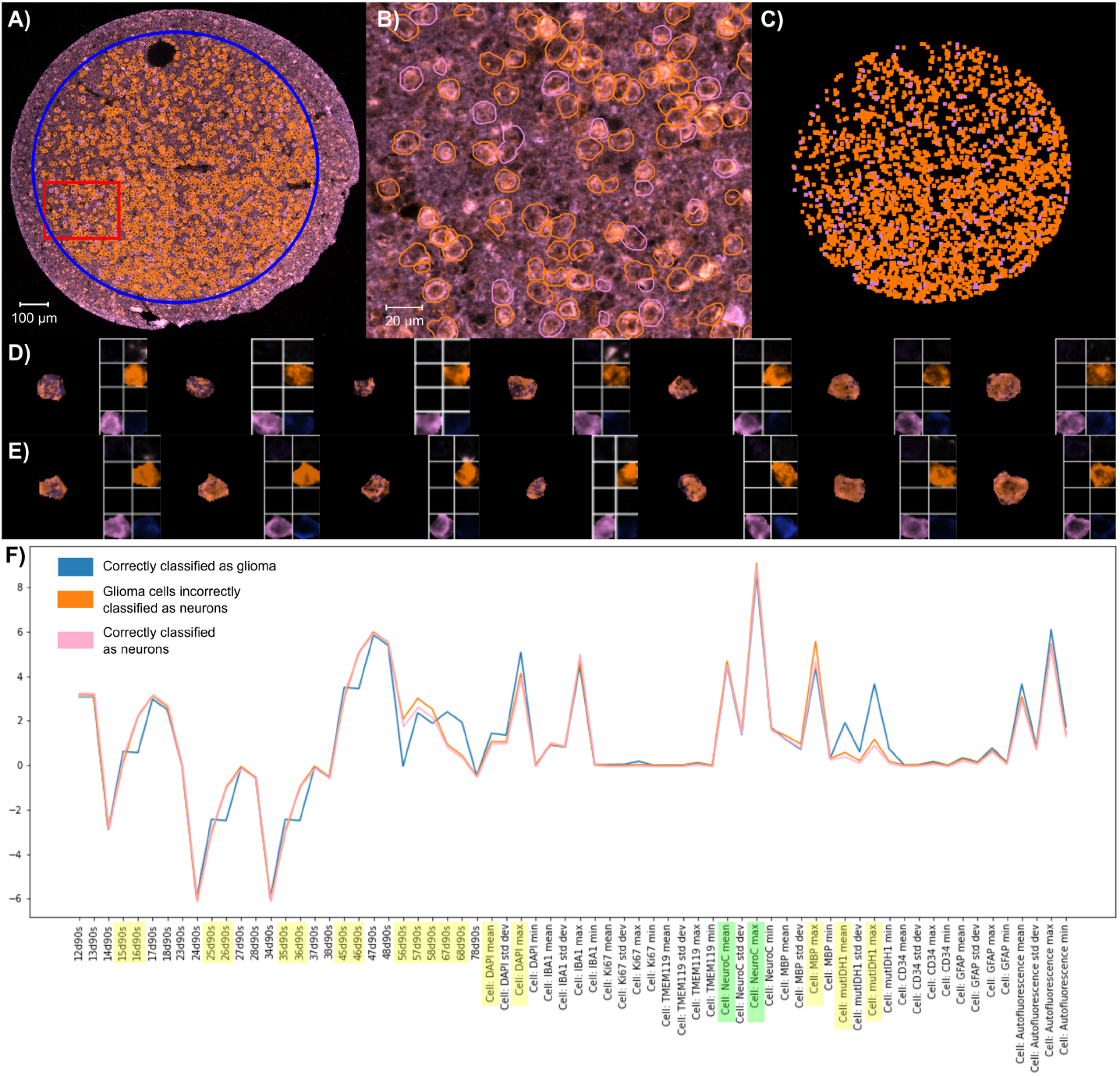
Illustration to explain confusion between glioma and neurons. A) shows the image data with channels for mutIDH1 and NeuroC. The blue circle shows the mask defining cells included in the enalysis. The core diameter is approximately 1.2 mm. B) shows a close up view of the core (scale bar 20um), where orange and pink outlines represent cells that were correctly (orange) or incorrectly (pink) classified by the FCNN as compared to expert 1. C) Plot showing the location of correctly classified glioma cells (orange) and glioma cells misclassified as neurons (pink). Note that the distribution of pink cells is homogeneous across the core therefore it is possible to conclude it is not an imaging artefact. D) Row displaying examples of incorrectly classified cells cells. E) Row displaying examples of correctly classified cells. All example cells in D and E contain miniviewers of each independent image channel. Note that both rows show presence of mutIDH1 and NeuroC at the same time. F) Visualization of feature profiles for the cell types. Blue and pink represent the average profile for all correctly classified cells of each class. Note that they are very similar and therefore difficult to distinguish. The profile in orange represents the incorrectly classified cells of this example core. Features highlighted in yellow contribute to the discriminative ability of the profiles, note that d90s are highlighted more. As expected mutIDH1 is an important factor. Features highlighted in green are related to NeuroC being a confusion factor. The values on the Y-axis are arbitrary units showing the relation between features and profiles.

**Figure S9:**
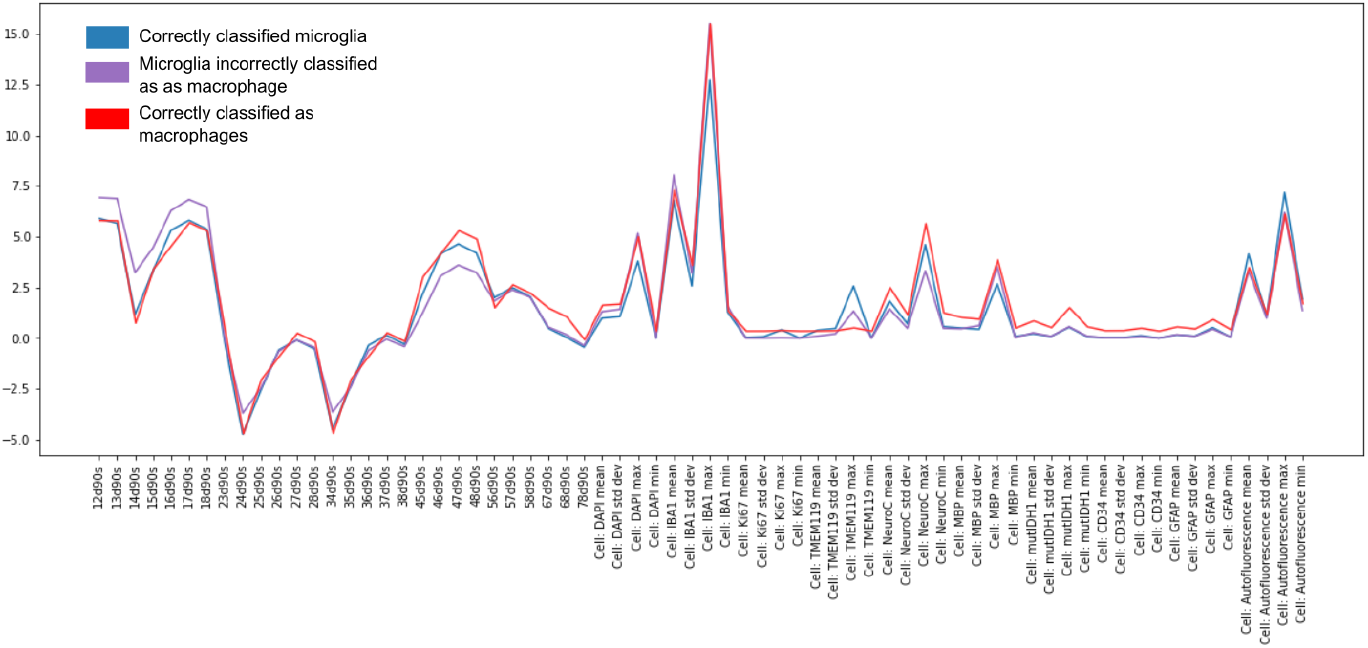
Illustration to explain confusion between celltypes.

**Figure S10:**
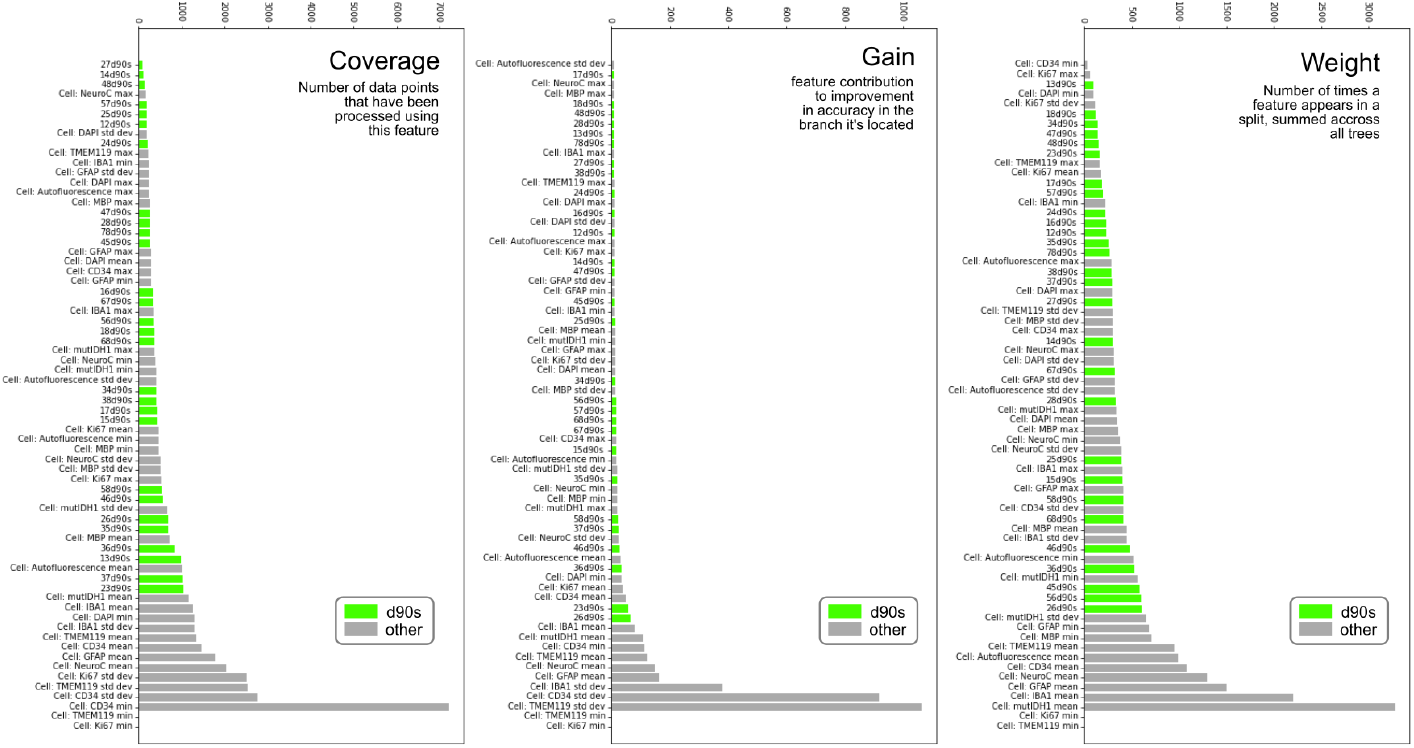
XGBoost measures of feature importance plotted for the average of each feature amongst all the trained 100 models. The importance of d90s can be observed in green, it is spread, not all d90s are equally important. Note that in all gain, weight and cover, the means of the markers have generally more importance than any other feature.

**Figure S11:**
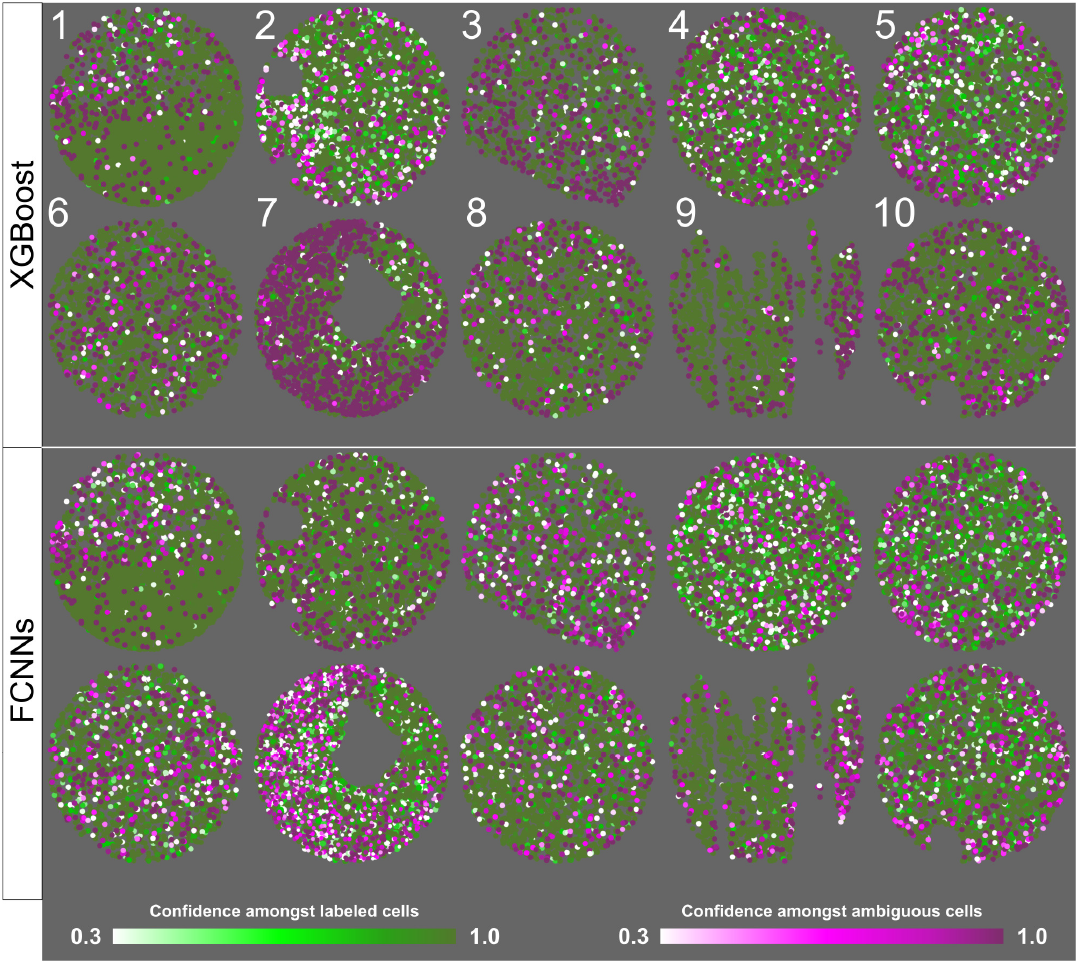
Colored confidence map for test cores. The color lookup table (LUT) is on purpose highlighting lower confidence cells. It can be observed confidence is spread across cores, for both methods. Green represents labeled cells by expert 1, and purple represents ambiguous cells according to expert 1. The diameter of each core is approximately 1.2 mm.

## Notes

https://www.lesliemachine.se/2021/04/24/ml-cell-class-neighborhood

### Competing Interest Statement

The authors have declared no competing interest.

### Summary of Updates

We have received wonderful, detailed and kind reviews which have helped us improve the manuscript. We have added attribution studies and further understanding of the sources of confusion of the models. We have also added further explanations on neighborhood studies

